# Regional deficits in endogenous regeneration of olfactory sensory neuron axons in the mouse olfactory bulb

**DOI:** 10.1101/2024.10.06.616907

**Authors:** Tenzin Kunkhyen, Kendall A. Curtis, Jordan D. Gregory, Thomas P. Deakin, Jane S. Huang, Claire E.J. Cheetham

## Abstract

Postnatal neurogenesis occurs in only a few regions of the mammalian nervous system. Hence, neurons that are lost due to neurodegenerative disease, stroke, traumatic brain injury or peripheral neuropathy cannot be replaced. Transplantation of stem cell-derived neurons provides a potential replacement strategy, but how these neurons can be encouraged to functionally integrate into circuits remains a significant challenge. In the mammalian olfactory epithelium (OE), olfactory sensory neurons (OSNs) continue to be generated throughout life from basal stem cells and can be repopulated even after complete ablation. However, the specialized population of navigator OSNs that ensures accurate olfactory receptor-specific targeting of OSN axons to glomeruli in the olfactory bulb (OB) is only present perinatally. Despite this, some studies have reported complete regeneration of specific glomeruli, while others have found various degrees of recovery, following OSN cell death. Variability in the extent of both initial OSN ablation and subsequent repopulation of the OE, and the focus on anatomical recovery, leave the extent to which newly generated OSNs can reinnervate the OB unclear. Here, we employed the olfactotoxic drug methimazole (MMZ) to selectively ablate OSNs without damaging the basal stem cells that generate them, enabling us to assess the extent of functional recovery of OSN input to the OB in the context of complete OSN repopulation. We found profound deficits in the recovery of odor-evoked responses in OSN axons in the glomerular layer of the dorsal OB five weeks after OSN ablation, a time point at which OSNs are known to have repopulated the OE. Histological analysis of mature OSN axons in the OB at 10 and 20 weeks post-methimazole showed persistent region-specific deficits in OSN axon reinnervation of the dorsal and medial OB, with the dorsomedial region being particularly adversely affected. In contrast, reinnervation of the glomerular layer in lateral and ventral regions of the OB was essentially complete by 10 weeks post-MMZ. Hence, we have identified a region-specific deficit in OSN reinnervation of the mouse OB, which sets the stage to identify the mechanisms that mediate successful vs. unsuccessful axonal regeneration in an endogenous population of stem cell-derived neurons.

## Introduction

Most mammalian neuronal types are generated only during embryonic and early postnatal development, meaning that in the adult nervous system, lost or damaged neurons cannot be replaced. In contrast, olfactory sensory neurons (OSNs) continue to be generated throughout life from basal stem cells in the olfactory epithelium (OE) of the nose of all terrestrial mammals [1–3]. OSNs therefore provide an opportunity to understand how an endogenous population of stem cell-derived neurons wires into highly organized neural circuits throughout life. Determining how OSNs achieve this, and particularly the extent to which OSN input to the brain can recover after substantial damage, will define mechanisms that may promote functional integration of transplanted stem cell-derived neurons to replace lost neurons elsewhere in the brain.

OSNs transmit odor input to the olfactory bulb (OB), with the axons of OSNs that express the same olfactory receptor coalescing together to form glomeruli [4]. Each mature OSN expresses a single olfactory receptor [5–7], which is expressed both on the cilia that protrude from the apical dendritic knob into the mucus layer on the surface of the OE, and on the axon, where it plays an important role in axon guidance [8–11]. The receptor therefore determines both the odor(s) that an OSN can bind and the glomerulus to which it projects its axon. Hence, there is a highly organized odor input map to the brain.

Globose basal cells (GBCs) generate newborn OSNs constitutively throughout life, with each OSN having a half-life of approximately one month [12–15]. In contrast, horizontal basal cells (HBCs) are normally quiescent and generate newborn OSNs only at key developmental time points and to enable repopulation following large-scale OSN cell death [16–18]. OSNs take 7-8 days from terminal cell division to reach maturity, which is defined by the onset of expression of the olfactory marker protein (OMP) [19–22]. During this time, in order to functionally integrate into the olfactory system, each OSN must select a single olfactory receptor, extend a dendrite to the apical surface of the OE, and project an axon to form synaptic connections with OB neuron dendrites in an appropriate glomerulus.

Anatomical recovery of the OSN population in the OE and of axonal projections to the OB has been studied using a variety of methods that induce OSN cell death, yielding mixed results. Olfactory nerve transection results in degeneration of OSN cell bodies and axons, but lost OSNs are rapidly replaced and reinnervate OB glomeruli within 2-5 weeks [23,24]. Similarly, the effects of treating the OE with Triton X-100 are reversible, with repopulation of the OE with OSNs and reinnervation of OB glomeruli occurring within 6 weeks, albeit with some disorganization of axonal projections [25]. Following inhalation of methyl bromide (MeBr) gas, most areas of the OE recover within six weeks although there is some persistent damage, and there is substantial reinnervation of OB glomeruli within 4-8 weeks, particularly in the anterior OB [26]. In contrast, treatment of the OE with zinc sulfate causes complete loss of the OE, resulting in long-term deficits in OSN repopulation and OB reinnervation even many months later [27]. More recently, systemic administration of methimazole (MMZ) has emerged as a popular model of OSN regeneration, as it avoids the confounds inherent to topical application, leaves the lamina propria intact [28], and spares basal cells such that OSNs can repopulate the OE in 4-12 weeks, depending on MMZ dosage [29–32].

Recovery of the olfactory receptor-specific topography of OSN projections to glomeruli has also received considerable attention. After selective ablation of nearly all OSNs expressing the P2 odorant receptor via a genetic approach, new P2-expressing OSNs accurately reinnervated the original P2 glomeruli in the OB [33]. In contrast, some degree of mis-targeting of OSN axons was seen after large-scale OSN cell death induced by several different methods. Following olfactory nerve transection, P2-expressing OSN axons projected to multiple widely distributed loci in the OB at 10 weeks, and while some further refinement occurred by 18 weeks, there was still a larger number of smaller P2 glomeruli than in control mice [34]. Similarly, 18 weeks after systemic dichlobenil treatment, a time point at which a significant number of P2-expressing OSNs have repopulated the OE, P2-expressing axons still terminated in numerous widely dispersed glomeruli [35]. After MeBr inhalation, 2A4(+) OSNs projected to 2-15-fold more glomeruli than in untreated mice, a projection pattern that was maintained up to five months post-treatment [36]. Following MMZ treatment, the accuracy of axonal targeting was better, but deficits remained. At three months post-MMZ, M72-expressing OSN axons largely projected to glomeruli in the expected dorsolateral location, but there were some wandering axons that were not pruned, as well as supernumerary M72 glomeruli [37]. Axonal projections of I7-expressing OSNs to ventral glomeruli largely recovered by 6 weeks post-MMZ, but formed two glomeruli within a small spatial region, rather than the expected one [37].

There are only three studies that have assessed functional recovery of odor input to the OB. The first study used *in vivo* synapto-pHluorin imaging of odor-evoked responses in the OB to directly assess functional recovery after MeBr treatment [38]. Substantial recovery of odor-evoked responses on the dorsal surface of the OB was seen, including reestablishment of odor-evoked response map topography. However, there was also evidence for atypical convergence of OSN axons, forming smaller glomeruli with less well-defined boundaries, especially when a high dose of MeBr was used [38]. Notably, OSN density did not recover fully after MeBr treatment, which may account for the deficits in OSN reinnervation of the OB [38]. The second study assessed recovery of behavioral performance six weeks after methimazole treatment and found significant recovery for a previously learnt odor discrimination task, but limited recovery of innate aversion to TMT [37]. The final study employed slice electrophysiology to show that functional reconnection of OSN axons with OB neurons occurs rapidly between 13-16 days after MMZ treatment, with presynaptic release probability already high, but some additional maturation occurring over the following week [29].

Therefore, important questions remain about the extent to which OSN axons can accurately reinnervate the OB to provide odor input to OB neurons. In this study, we employed MMZ treatment to address these questions, because it does not damage the lamina propria or basal stem cell population, enabling complete OSN repopulation of the OE. We found profound deficits in OSN reinnervation of the dorsal region of the OB, with the dorsomedial region recovering particularly poorly. Recovery of odor-evoked responses in the dorsal OB was very limited, and OMP immunohistochemistry in fixed tissue confirmed deficits in reinnervation of dorsal, and to a lesser extent medial, glomeruli at 10 weeks post-MMZ. Some of these deficits persisted at 20 weeks post-MMZ, while others recovered at this later time point. In contrast, we did not detect any differences between lateral and ventral OB glomeruli in MMZ-treated vs. control mice.

## Methods

### Experimental animals

Animal procedures conformed to National Institutes of Health and ARRIVE guidelines and were approved by the University of Pittsburgh Institutional Animal Care and Use Committee. Mice were maintained on a 12 h light/dark cycle in individually ventilated cages at 22 °C and 48% humidity with unrestricted access to food and water unless otherwise stated. Mice were group-housed if same sex littermates were available.

C57BL/6J mice (strain #000664) were purchased from The Jackson Laboratory while other lines were bred in-house. Generation of the OMP-IRES-tTA [39] (Jax #017754), tetO-GCaMP6s [40] (Jax #024742), TAAR4-RFP [41] and TAAR4-ChR2-YFP [41] lines has been published. TAAR4-RFP and TAAR4-ChR2-YFP mice were obtained from Dr. Thomas Bozza (Northwestern University). Each experimental group comprised approximately equal numbers of male and female mice. Mice were genotyped by PCR using previously validated primers [40–45]. Experimental mice were C57BL/6J, OMP-GCaMP6s [OMP-IRES-tTA^+/−^;tetO-GCaMP6s^+/−^], TAAR4-RFP homozygotes or TAAR4-ChR2-YFP homozygotes. For TAAR4 analysis, the saline group consisted of one TAAR4-RFP and two TAAR4-ChR2-YFP mice, while the MMZ group consisted of three TAAR4-RFP mice. A total of 37 mice were used in this study.

### MMZ ablation of OSNs

Mice received a single intraperitoneal injection of 75 mg/kg MMZ dissolved in sterile saline. For chronic imaging and for analysis of TAAR4-RFP and TAAR4-ChR2-YFP mice, an equivalent volume of sterile saline was administered to control mice. For OMP immunohistochemistry, control mice consisted of a combination of saline-injected mice (n = 2 per time point) and age matched (18- or 28-week-old) C57BL/6J mice that were purchased as controls and did not receive saline (n = 3 per time point).

### Cranial window implantation and headbar attachment

8-10-week-old OMP-GCaMP6s mice were anesthetized with isoflurane (4% for induction, 1.5-2% for maintenance, in 1l/min O_2_) and received 5 mg/kg ketoprofen. A 3 mm diameter craniotomy over both OBs was made and a 3 mm diameter glass coverslip was implanted using superglue and dental cement before re-suturing the scalp, as described in detail [46]. Two weeks later, the scalp was removed, and a head bar was attached to enable reproducible repositioning under the 2-photon microscope.

### Chronic *in vivo* 2-photon imaging and image analysis

Mice were lightly anesthetized with 5 mg/kg dexmedetomidine and 1% sevoflurane. Chronic *in vivo* 2-photon imaging was performed using a Bergamo II 2-photon microscope (ThorLabs) equipped with a SemiApo 20x/1.0NA water-immersion objective (Olympus) and an Insight X3 IR laser (Newport) mode-locked at 935 nm. Time-lapse images were collected at 15 fps and were 656 μm^2^ (pixel size 1.28 μm). Odorant stimulation was performed using a custom-built Arduino-controlled olfactometer. Stimulation and image acquisition timing were controlled and recorded using a custom-written Python-based user interface, ThorSync and ThorImage 4.0 or 4.1 (ThorLabs). The odorant panel consisted of seven odorants known to activate the dorsal surface of the OB: ethyl butyrate, hexanal, 2-hexanone, propionic acid, isoamyl acetate, methyl salicylate and acetophenone (all 1% v/v dilutions in mineral oil). Saturated vapor from odorant vials, or a blank stimulus consisting of deodorized dehumidified air, were delivered into the anesthetic carrier stream at a constant flow rate of 1 l/min. Odor triggering and delivery were checked before each imaging session. Because we expected no or minimal odorant-evoked responses at early time points post-MMZ, we designed our experiments such that we always imaged a saline-injected mouse on the same day as an MMZ-injected mouse. This ensured that an absence of odorant-evoked responses in an MMZ-injected mouse could be attributed to the MMZ treatment rather than any technical issues.

Image trials were 20 s in duration, consisting of 4s baseline, a 1s stimulus and 15 s post-stimulus. Three trials were delivered for each stimulus, in a pseudorandomized order with 90 s between trials. Data was collected for two to three fields of view (FOVs) per mouse. For each FOV, the xyz co-ordinates of the imaged location were recorded, and a widefield camera image of the surface vasculature was captured, to enable the same locations to be re-imaged at subsequent time points. One week later, each mouse was anesthetized and the same FOVs were identified using the XY co-ordinates of the microscope system, the pattern of surface vasculature and the pattern of baseline GCaMP6s fluorescence in glomeruli, when present (note that this latter parameter was less useful in MMZ-treated mice due to gradual degeneration of existing OSN axons). Odor-evoked responses were collected as for the first imaging session. Mice received either MMZ or saline immediately after the second imaging session. Windows were inspected each week for the presence of any bone regrowth that could occlude the imaged FOVs, and imaging was repeated every week for 10 weeks if windows remained clear. Only mice with at least 7 weeks of imaging sessions were included in the analysis. Imaging occurred discontinuously for longer time periods (up to 21 weeks) for some mice in which windows remained clear.

Images were analyzed using two approaches, both in Fiji [47]. In the glomerulus-based approach, individual glomeruli were outlined in the week 1 images. For each stimulus, the three trials were averaged, and a maximum intensity projection (MIP) was generated. Outlines of all visible glomeruli were drawn for the first stimulus MIP using the polygon tool, superimposed on each subsequent stimulus MIP, and additional visible glomeruli were outlined until all eight MIPs had been completed, to generate the final set of glomeruli for analysis. The unbiased grid-based approach was designed to detect any recovery of odor-evoked responses in MMZ-treated mice, including in locations that did not previously respond and so could be missed using the glomerulus-based approach. A 16-square grid (20.5 μm per square) was overlaid on each image irrespective of glomerular position. Mean fluorescence intensity was then extracted for each glomerulus or grid square for each trial. All images were analyzed using both approaches.

### Perfusion, immunohistochemistry, widefield microscopy and image analysis

Perfusion, OB dissection, cryopreservation and sectioning were performed as described [48]. For OMP immunohistochemistry, three 40 μm coronal OB sections per mouse at 25%, 50% and 75% along the anterior-posterior axis were stained to detect OMP using anti-OMP primary antibody (#544-10001, Wako Chemicals; 1:5000, 96 h at 4 °C) and donkey anti-goat-Alexa Fluor 546 (AF546) secondary antibody (1:500, 1 h at room temperature). For TAAR4 glomerulus analysis, 40 μm coronal sections from 1.0-1.8 mm along the anterior-posterior axis of the OB were collected and stained with either anti-RFP-ATTO594 nanobody (rba594, Chromotek; for sections from TAAR4-RFP mice) or anti-GFP-ATTO488 nanobody (gba488, Chromotek; for sections from TAAR4-ChR2-YFP mice) at 1:500 for 1 hour at room temperature. All sections were mounted using Vectashield containing DAPI. Tiled images of DAPI and OMP signals for OMP staining, or DAPI and RFP or YFP staining for TAAR4-labled mice, throughout entire OB sections were acquired using a Revolve microscope with Echo software (Echo) equipped with a 10x Plan Apo 0.4 NA air objective (Olympus). For analysis of TAAR4 glomeruli, all stained sections were visually surveyed and tiled images of the dorsal half of any sections containing RFP+ or YFP+ axons were collected. Bandpass excitation and emission filter wavelengths were (in nm): DAPI (385/30, 450/50), YFP-ATTO488 (470/40, 525/50) and OMP-AF546 or RFP-ATTO594 (530/40, 590/50). Images were stitched using Affinity Photo software (Affinity) to generate a complete image of each section.

Image analysis was performed using Fiji (ImageJ) [47]. For analysis of OMP-stained sections, the glomerular layer (GL) was outlined in the DAPI channel of each section and subdivided into dorsomedial, dorsolateral, medial, ventral and lateral regions. This subdivision of the dorsal GL into dorsomedial and dorsolateral regions was based on initial qualitative observations that OMP staining was not uniform throughout the dorsal GL in MMZ-treated mice. The area of each GL region was determined. Individual glomeruli were also outlined in the DAPI channel, with outlines verified and edited in the OMP channel. The number of glomeruli in each GL region was counted, using both the DAPI and OMP channels to ensure that all glomeruli were counted. Glomerular density was defined as the number of glomeruli within a GL region divided by the area of that region. The perimeter and circularity [4π(area/perimeter^2^), where a value of 1.0 indicates a perfect circle], of each glomerulus were also determined. The percentage of the GL occupied by glomeruli was determined for each GL region using the total region area and the summed area of all glomeruli therein. We then calculated the background fluorescence, using 300-pixel diameter circles located in the dorsal, medial, ventral and lateral external plexiform layer. This background fluorescence value was multiplied by two and used to threshold the OMP channel to determine the area fraction, i.e. the percentage of each glomerulus filled by OMP+ pixels. For analysis of TAAR4 glomeruli, we identified the section containing the largest cross-section of each TAAR4 glomerulus and outlined the glomerulus using the DAPI and RFP/YFP channels. Perimeter and area fraction were determined as for OMP-stained sections, using RFP or YFP fluorescence to determine percentage filling of the TAAR4 glomerulus by TAAR4+ axons.

### Statistics

Data were analyzed using Prism 10 (GraphPad) with alpha = 0.05. For comparison of glomerulus- and grid-based OMP-GCaMP6s values, data passed the Shapiro–Wilk test for normality and were analyzed using one- or two-way ANOVA. For analysis of OMP-stained OB sections, data passed the Shapiro–Wilk test for normality and were analyzed using two-way ANOVA with Holm-Sidak’s multiple comparisons tests between 10-week MMZ vs. control and 20-week MMZ vs. control groups. For comparison of TAAR4 glomeruli from MMZ- vs. saline-treated mice, data passed the Shapiro–Wilk test for normality and were analyzed using Welch’s unpaired t-tests. The details of all statistical tests and their outcomes are provided in the figure legends.

We employed generalized linear models (GLMs) implemented in R (R Project for Statistical Computing, https://www.r-project.org) to assess the effect of position along the anterior-posterior axis and interactions between anterior-posterior axis position and MMZ treatment. GLMs were of the form g [E(Y)] = a + β_1_(MMZ) + β_2_(anterior) + β_3_(posterior) + β_4_(DM) + β_5_(lateral) + β_6_(medial) + β_7_(ventral) + β_8_(time) + β_9_(MMZ:time) + β_10_(MMZ:anterior) + β_11_(MMZ:posterior) + β_12_(MMZ:DM) + β_13_(MMZ:lateral) + β_14_(MMZ:medial) + β_15_(MMZ:ventral), where g is the link function; E(Y) is the expected value of the response variable, Y; MMZ, anterior, posterior, DM, lateral, medial, ventral and time are explanatory variables with : denoting interactions between explanatory variables, and a and β_1_ …β_15_ are coefficients of the model. MMZ treatment was modeled with a dummy explanatory variable, MMZ, for which 1 denoted MMZ-treated mice and 0 denoted control mice. Position along the anterior-posterior axis was modeled with two dummy explanatory variables for which anterior (anterior section 1, otherwise 0) denoted sections 25% along the anterior-posterior axis and posterior (posterior section 1, otherwise 0) denoted sections 75% along the anterior-posterior axis, such that data from central sections are encoded (anterior, posterior) by (0,0), data from anterior sections are encoded by (1,0) and data from posterior sections are encoded by (0,1). GL region was modeled with four dummy explanatory variables in which DM (DM GL 1, otherwise 0) denoted the DM GL, lateral (lateral GL 1, otherwise 0) denoted the lateral GL, medial (medial GL 1, otherwise 0) and ventral (ventral GL 1, otherwise 0) denoted the ventral GL. As such, data from the DL GL region are encoded (DM, lateral, medial, ventral) by (0,0,0,0), data from the DM GL region are encoded by (1,0,0,0), data from the lateral GL region are encoded by (0,1,0,0), data from the medial GL region are encoded by (0,0,1,0) and data from the ventral GL region are encoded by (0,0,0,1). Experimental time point was modeled with a dummy explanatory variable, Time, for which 1 denoted the 20-week time point and 0 denoted the 10-week time point. For each response variable (glomerular density, % GL fill by glomeruli, glomerular perimeter, % glomerular fill by OMP+ OSN axons) we tested gaussian and Gamma error distribution families and identity, inverse and log link functions. The best model for the data was found by minimizing Akaike’s information criterion for nested models.

## Results

### Poor recovery of odorant-evoked responses in dorsal OB glomeruli after MMZ treatment

To determine the timeline and extent of recovery of odor-evoked responses in mature OSN axons in the OB after MMZ treatment, we implanted adult OMP-GCaMP6s mice with OB cranial windows (Fig. 1a), followed by head bars to enable reproducible repositioning under the 2-photon microscope. Three weeks after window implantation, we performed a baseline 2-photon imaging session of responses evoked by a panel of seven odorants known to activate glomeruli on the dorsal surface of the OB. A second baseline imaging session was performed one week later, immediately after which mice were treated with either 75 mg/kg MMZ or saline as a vehicle control. Subsequent imaging sessions were performed weekly for at least five weeks post-treatment (Fig. 1b,c).

**Figure 1.**
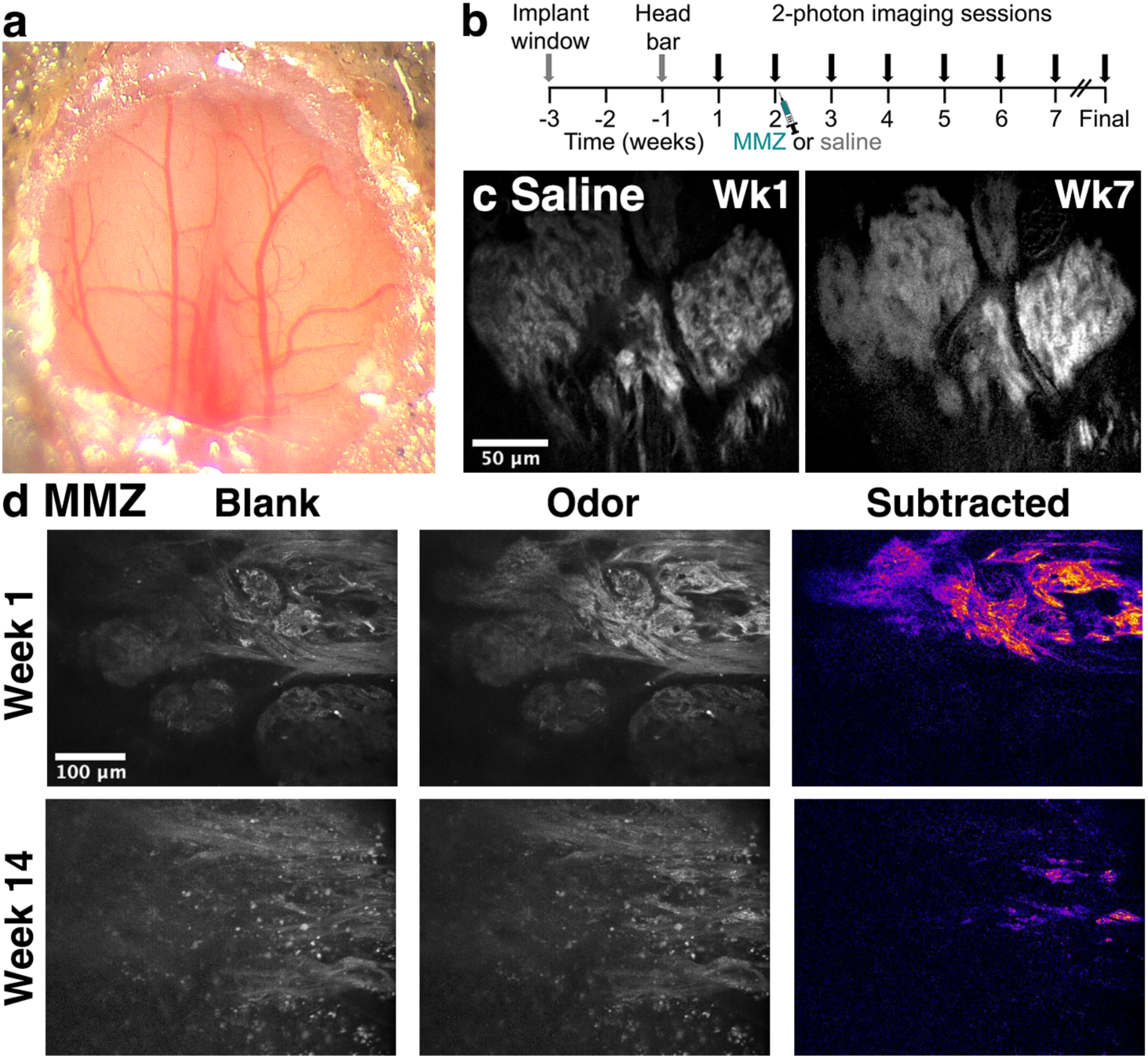
Chronic imaging of odorant-evoked responses in OMP-GCaMP6s mice. **a.** 3 mm diameter chronic cranial window over both OBs. **b.** Timeline for chronic imaging of OMP-GCaMP6s mice. **c.** 2-photon images showing stability of odorant-evoked responses across time in saline-injected control mice. Images are blank-subtracted responses to 1% ethyl butyrate. **d.** 2-photon images showing poor recovery of odorant-evoked responses 12 weeks after MMZ treatment. Responses to blank and odor (1% ethyl butyrate) are shown, with the blank-subtracted odor-evoked responses shown on a fire lookup table to highlight the weak responses post-MMZ.

Analysis of chronic functional imaging data in the OB has typically involved comparison of odorant response amplitudes within individual glomeruli over time [38,49–51]. However, we reasoned that regeneration of OSN input to the OB might occur outside the glomerular regions that were defined prior to MMZ treatment based on their baseline GCaMP6s fluorescence and/or responsiveness to one of the tested odorants. Therefore, we developed an unbiased grid-based approach to measure odorant-evoked responses in the same fields of view over time (Fig. 2a). We validated this grid-based approach by comparison to the data from glomerulus-based analysis using data from saline-treated mice. The amplitude of responses varied somewhat over time but was broadly similar across seven weeks of imaging (Fig. 2b), similar to a previous study [38]. The amplitudes of odorant-evoked responses were consistently larger in glomeruli than in grid squares (Fig. 2c), as would be expected given the inclusion of non-odorant-responsive areas within grid squares. However, the ratio of glomerulus to grid responses amplitudes was similar across odorants (Fig. 2d). Therefore, we concluded that the grid-based approach could be used to detect recovery of odorant-evoked responses anywhere in the imaged fields of view in MMZ-treated mice.

**Figure 2.**
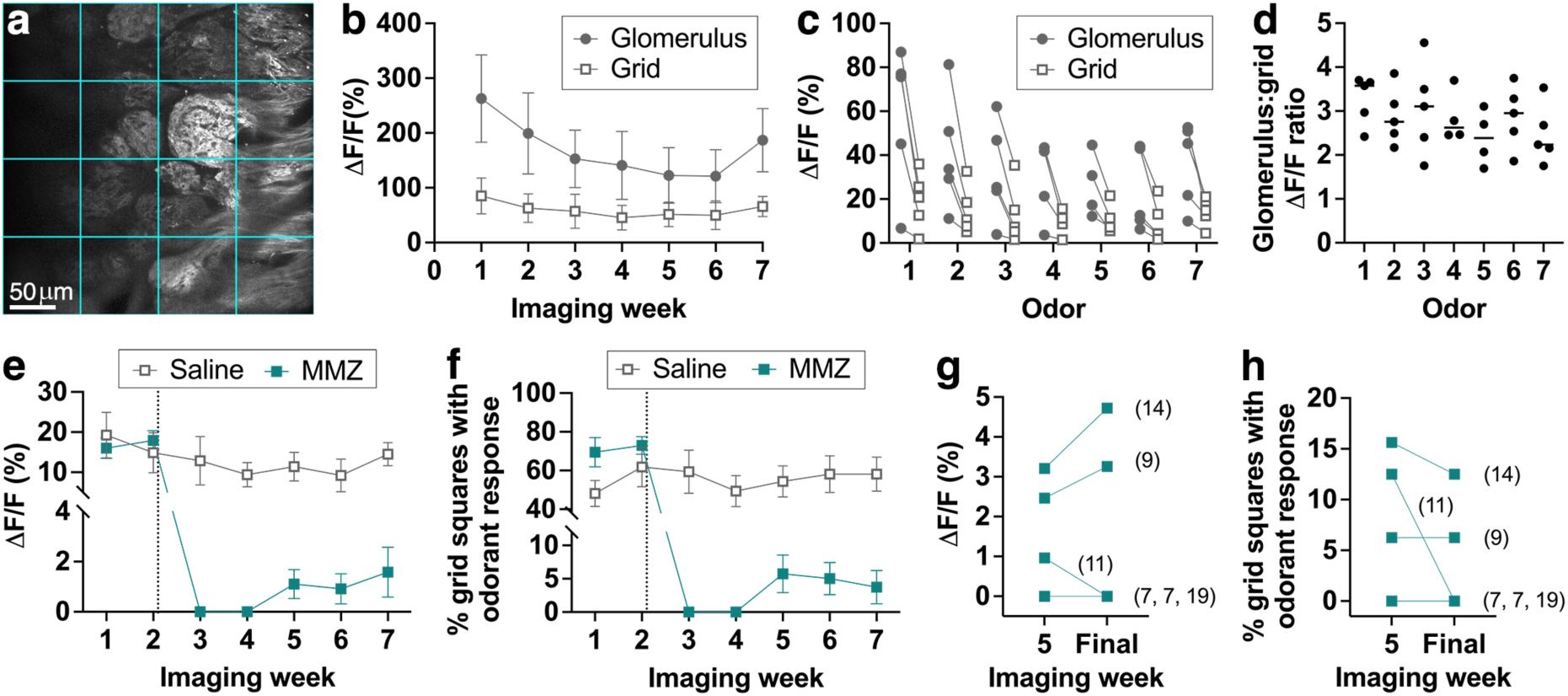
Poor recovery of odorant-evoked responses in dorsal OB glomeruli 10-20 weeks after MMZ treatment. **a.** Schematic of grid-based analysis approach. **b.** Stability of ethyl butyrate-evoked response amplitude in saline-treated OMP-GCaMP6s mice across 7 weeks of chronic imaging using glomerulus-based or grid-based analysis. Data are shown as mean ± s.e.m., n = 5 mice. **c.** Odorant-evoked response amplitude is consistently smaller across odorants for the grid-based approach. Lines connect data points for individual mice. P = 0.094, F_2,16_ = 2.73 for effect of odorant; P = 0.014, F_1,8_ = 9.67 for glomerulus vs. grid-based analysis; P = 0.57, F_6,44_ = 0.81 for interaction; two-way ANOVA. n = 4-5 mice per odorant. Note that data is not shown for mouse-odorant pairs in which no response was detected using either approach. **d.** Ratio of odorant-evoked response amplitude for the glomerulus-based vs. grid-based approaches is similar across odorants. P = 0.55, F_6,26_ = 0.84, one-way ANOVA. n as in c. **e.** Effect of MMZ on mean response amplitude per grid square across all tested odorants. Dotted line indicates injection of saline or MMZ after the second imaging session. n = 5 saline- and 6 MMZ-treated mice. **f.** Effect of MMZ on percentage of grid squares that responds to any of the tested odorants. Dotted line and n as in e. **g,h.** No further recovery of odorant-evoked responses detected in grid squares between imaging week 5 and the final time point that could be imaged in each mouse. Numbers in parentheses indicate number of weeks for which imaging was performed. n = 6 MMZ-treated mice. **g.** Mean response amplitude across all odorants. **h.** Percentage of grid squares that responds to any of the tested odorants.

We next used our grid-based approach to assess recovery of odorant-evoked responses in MMZ-treated mice. Responses in weeks 1 and 2, prior to MMZ treatment, were similar, with no difference in odorant-evoked response amplitude or the percentage of grid squares that were activated by at least one odorant (Fig. 2e,f). One week after MMZ treatment (imaging week 3), odorant-evoked responses were completely absent (Fig. 2e,f; imaging week 3), confirming that MMZ had resulted in OSN ablation. Odorant-evoked responses remained completely absent at two weeks post-MMZ (imaging week 4), but by three weeks post-MMZ (imaging week 5), small areas of odorant-responsive axons were present in some mice. Surprisingly, neither the amplitude of odorant-evoked responses (Fig. 2e), nor the percentage of grid squares responding to at least one odorant (Fig. 2f) increased further at five weeks post-MMZ (imaging week 7), a time point at which the OE has been repopulated with OSNs [30,32]. Indeed, in a subset of mice in which we performed additional imaging sessions over longer time periods, even up to 17 weeks post-MMZ (imaging week 19), we observed no further recovery of odorant-evoked responses beyond that seen at 3 weeks post-MMZ (imaging week 5; Fig. 2g,h). Overall, we concluded that recovery of odorant-evoked responses in the dorsal OB following MMZ treatment is limited, and indicates a deficit in mature OSN reinnervation of the dorsal OB.

### Region-specific deficits in OSN reinnervation 10 and 20 weeks after MMZ treatment

To determine whether the deficits in OSN reinnervation suggested by our *in vivo* calcium imaging data were specific to the dorsal surface of the OB, we employed an immunohistochemical approach that enabled us to analyze entire OB sections. 8-week-old C57BL/6J mice were injected with MMZ and perfused 10 or 20 weeks later, to allow for the possibility that further regeneration might occur over longer time periods than were consistently feasible to assess using our *in vivo* imaging approach. Age-matched control C57BL/6J mice were also perfused, and OB sections from MMZ-treated and control mice were stained for OMP to detect mature OSN axons.

We observed that OMP staining appeared similar in the medial, lateral and ventral regions of the GL in control and MMZ-treated mice. In contrast, OMP staining was less extensive on the dorsal surface of the OB of MMZ-treated mice compared to controls, with fewer glomeruli present and some glomeruli lacking innervation by OMP+ OSNs (Fig, 3). It was also apparent that these differences were not uniform throughout the dorsal region of the OB (Fig. 3). Therefore, we subdivided the GL into dorsomedial (DM), dorsolateral (DL), medial, lateral and ventral regions for quantitative analysis (Fig. 4a).

**Figure 3.**
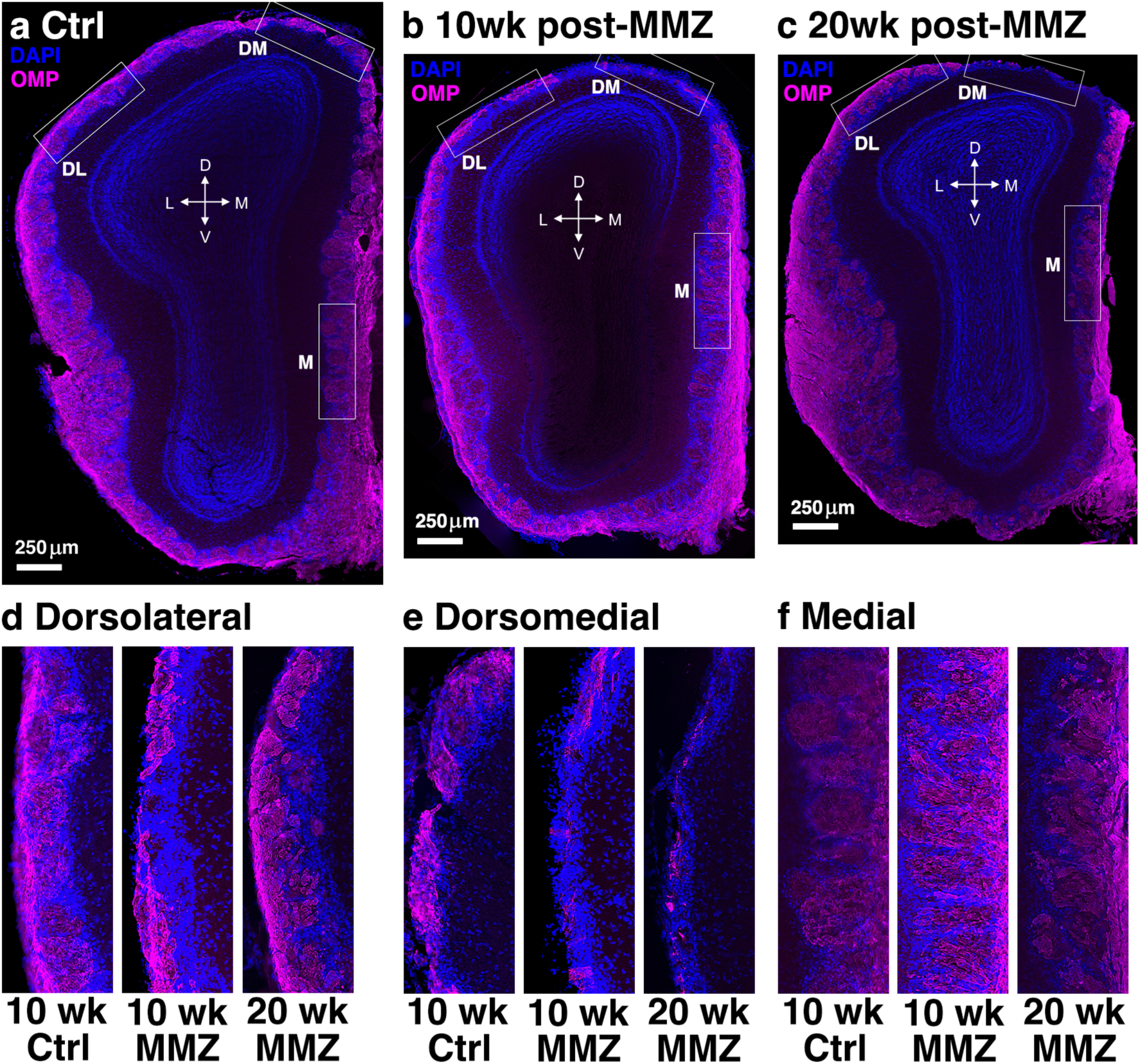
OMP staining reveals region-specific deficits in OSN reinnervation of the OB up to 20 weeks post-MMZ. Example widefield images of coronal OB sections stained for OMP and DAPI. **a.** Control mouse age-matched to the 10-week MMZ time point. **b.** 10-week post-MMZ mouse. **c.** 20-week post-MMZ mouse. **d.** Enlarged view of dorsolateral regions shown by boxes in a-c. **e.** Enlarged view of dorsomedial regions shown by boxes in a-c. **f.** Enlarged view of medial regions shown by boxes in a-c.

**Figure 4.**
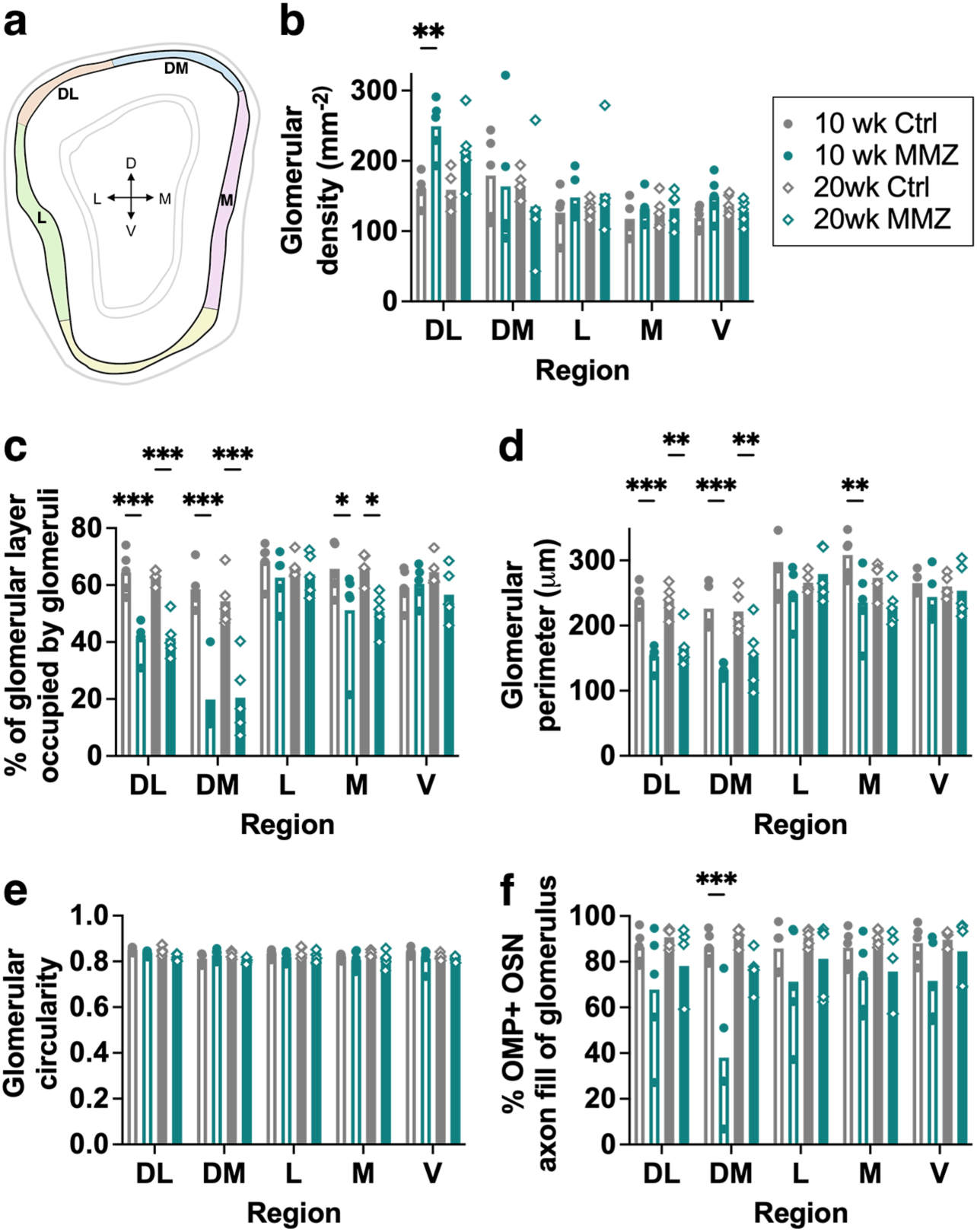
Region-specific deficits in regeneration of OSN axons 10-20 weeks after MMZ treatment. **a.** Schematic showing GL regions that were analyzed. Arrows show dorsal (D), ventral (V), medial (M) and lateral (L) directions. Analyzed regions are dorsolateral (orange), dorsomedial (blue), lateral (green), medial (pink), and ventral (yellow). **b.** Glomerular density is higher in the DL region of the GL 10 weeks post-MMZ. P < 0.001, F_4,80_ = 8.50 for GL region; P = 0.099, F_3,80_ = 2.16 for effect of MMZ; P = 0.23, F_12,80_ = 1.32 for interaction, 2-way ANOVA with Holm-Sidak multiple comparisons. For 10-week control vs. MMZ: DL P = 0.007, t = 3.33; DM P = 0.91, t = 0.57; lateral P = 0.94, t = 0.81; medial P = 0.99, t = 0.54; ventral P = 0.84, t = 1.13. For 20-week control vs. MMZ: DL P = 0.15, t = 2.08; DM P = 0.82, t = 1.04; lateral P = 0.94, t = 0.76; medial P = 1.00, t = 0.21; ventral P = 0.97, t = 0.40. **c.** Percentage of the GL occupied by glomeruli is reduced in the DM, DL and medial regions 10 and 20 weeks post-MMZ. P < 0.001, F_4,80_ = 30.4 for GL region; P < 0.001, F_3,80_ = 31.1 for effect of MMZ; P < 0.001, F_12,80_ = 4.63 for interaction, 2-way ANOVA with Holm-Sidak multiple comparisons. For 10-week control vs. MMZ: DL P < 0.001, t = 4.39; DM P < 0.001, t = 7.27; lateral P = 0.86, t = 1.09; medial P = 0.028, t = 2.72; ventral P = 0.91, t = 0.23. For 20-week control vs. MMZ: DL P < 0.001, t = 6.35; DM P < 0.001, t = 6.35; lateral P = 0.95, t = 0.48; medial P = 0.026, t = 2.94; ventral P = 0.60, t = 1.48. **d.** Region-specific reduction in glomerular perimeter after MMZ treatment. P < 0.001, F_4,80_ = 28.8 for GL region; P < 0.001, F_3,80_ = 19.8 for effect of MMZ; P = 0.046, F_12,80_ = 1.90 for interaction, 2-way ANOVA with Holm-Sidak multiple comparisons. For 10-week control vs. MMZ: DL P < 0.001, t = 4.06; DM P < 0.001, t = 4.40; lateral P = 0.14, t = 2.28; medial P = 0.005, t = 3.43; ventral P = 0.90, t = 0.99. For 20-week control vs. MMZ: DL P = 0.004, t = 3.37; DM P = 0.006, t = 3.20; lateral P = 0.78, t = 0.65; medial P = 0.18, t = 2.01; ventral P = 0.97, t = 0.29. **e.** Glomerular circularity is similar in MMZ-treated and control mice. P = 0.012, F_4,80_ = 3.47 for GL region; P = 0.005 F_3,80_ = 4.67 for effect of MMZ; P = 0.81, F_12,48_ = 0.62 for interaction, 2-way ANOVA with Holm-Sidak multiple comparisons. For 10-week control vs. MMZ: DL P = 0.71, t = 0.93; DM P = 0.77, t = 1.03; lateral P = 0.75, t = 0.98; medial P = 0.75, t = 1.05; ventral P = 0.36, t = 1.75. For 20-week control vs. MMZ: DL P = 0.14, t = 3.30; DM P = 0.49, t = 1.60; lateral P = 0.69, t = 1.37; medial P = 0.53, t = 1.49; ventral P = 0.93, t = 0.53. **f.** Selective deficit in innervation of DM glomeruli by OMP+ OSN axons. P = 0.17, F_4,80_ = 2.30 for GL region; P < 0.001, F_3,80_ = 16.1 for effect of MMZ; P = 0.20, F_12,80_ = 1.35 for interaction. 2-way ANOVA with Holm-Sidak multiple comparisons. For 10-week control vs. MMZ: DL P = 0.20, t = 2.05; DM P < 0.001, t = 5.35; lateral P = 0.45, t = 1.60; medial P = 0.51, t = 1.41; ventral P = 0.31, t = 1.82. For 20-week control vs. MMZ: DL P = 0.52, t = 1.39; DM P = 0.30, t = 1.62; lateral P = 0.72, t = 1.03; medial P = 0.44, t = 1.62; ventral P = 0.92, t = 0.56. n = 5 mice per group for all analyses.

We first determined whether the density of glomeruli within each region of the GL was affected at 10 or 20 weeks post-MMZ treatment. Glomerular density was similar throughout much of the GL; however, compared to controls, glomerular density was significantly higher in the DL region 10 weeks post-MMZ (Fig. 4b). By 20 weeks post-MMZ, there remained a trend towards higher glomerular density in the DL region, but there was no significant difference compared to controls (Fig. 4b), suggesting some gradual ongoing regeneration. Glomerular density was not dependent on position along the anterior-posterior axis, nor was there an interaction between anterior-posterior axis position and the effect of MMZ treatment (Table S1).

We reasoned that the increase in DL glomerular density could be due to a larger number of glomeruli and/or a smaller DL GL area. Furthermore, the lack of a difference in glomerular density elsewhere in the GL was somewhat surprising based on visual observation of OB sections (Fig. 3). However, if the area of the GL was reduced by MMZ treatment in other regions, there could be fewer glomeruli but no change in glomerular density. Therefore, we next determined the percentage of the GL that was occupied by glomeruli for each region. Indeed, we found that the percentage of the GL occupied by glomeruli was significantly lower in the DL and DM regions at both 10 and 20 weeks post-MMZ, and in the medial region at 10 weeks post-MMZ (Fig. 4c). The magnitude of this reduction was greatest in the DM region and smallest in the medial region (DL 10 week −36%; DL 20 week −34%; DM 10 week −66%; DM 20 week: −62%; medial 10 week −22%; percentage changes in mean values, Fig. 4c). In contrast, there was no difference in the percentage of the GL occupied by glomeruli in the lateral or ventral regions between MMZ-treated and control mice at either time point (Fig. 4c). There was also no effect of position along the anterior-posterior axis on the GL percentage occupied by glomeruli, and no interaction between anterior-posterior axis position and MMZ treatment that affected GL filling by glomeruli (Table S1). Therefore, we concluded that MMZ treatment causes a persistent reduction in the number of glomeruli in the dorsal and medial GL, with this effect being particularly pronounced in the DM region.

We next analyzed the impact of MMZ treatment on the glomeruli themselves. We first determined the impact of MMZ treatment on glomerular size across GL regions. At 10 weeks post-MMZ, glomerular perimeter was significantly smaller in the DL, DM and medial regions compared to controls, whereas glomerular perimeter was unaffected in the lateral and ventral GL (Fig. 4d). By 20 weeks post-MMZ, glomerular perimeter remained significantly smaller in the DL and DM regions, but there was no longer a difference in the medial region, with the lateral and ventral GL again unaffected (Fig. 4d). We also found no effect of anterior-posterior axis position on glomerular perimeter and no interaction between anterior-axis position and MMZ treatment that affected glomerular perimeter (Table S1). We then analyzed the circularity of glomeruli to determine whether the shape of glomeruli that regenerated following MMZ treatment differed from glomeruli in control mice. Glomerular circularity was unaffected by MMZ treatment (Fig. 4e). We therefore concluded that MMZ treatment results in smaller glomeruli throughout the dorsal and medial GL, but this effect is more persistent in the dorsal OB. In contrast, glomerular shape was minimally impacted by MMZ treatment.

The final parameter that we analyzed was the percentage of individual glomeruli that was occupied by OMP+ OSN axons, to provide a measure of glomerular filling by mature OSN axons. We found that in the DM region, the percentage of a glomerulus that was filled by OMP+ axons was strongly and significantly reduced at 10 weeks post-MMZ (mean reduction 56%, Fig. 4f). By 20 weeks, there was no longer a statistically significant difference in the DM region (mean reduction 16%, Fig. 4f), suggesting ongoing recovery between 10 and 20 weeks post-MMZ. There was no difference in mature OSN axon filling of glomeruli between control and MMZ-treated mice for the DL, lateral, medial or ventral regions of the GL. The position of analyzed sections along the anterior-posterior axis also did not affect glomerular filling by OMP+ OSN axons and there was no interaction between anterior-posterior axis and the effect of MMZ treatment that affected OMP+ OSN axon filling of glomeruli (Table S1). Hence, we concluded that there was a selective deficit in reinnervation of DM region glomeruli by mature OSNs following MMZ treatment that gradually recovers over long time periods.

### Deficits in OSN reinnervation of TAAR4 glomeruli after MMZ treatment

Our analysis of OMP staining did not allow us to directly compare the same glomeruli between MMZ-treated and control mice. Therefore, we used fluorescently tagged mouse lines to compare the size and OSN filling of TAAR4 glomeruli, which are found in the DM OB, 10 weeks after MMZ or saline treatment. We identified 7 TAAR4 glomeruli in sections from three saline-treated mice (Fig. 5a), and 8 TAAR4 glomeruli in sections from three MMZ-treated mice (Fig. 5b) based on clear coalescence of TAAR4+ OSN axons, suggesting qualitatively that TAAR4 glomeruli may be amongst the relatively rare DM glomeruli that exhibit significant reinnervation following MMZ treatment. We found no effect of MMZ treatment on TAAR4 glomerulus perimeter (Fig. 5c), in agreement with our analysis of the larger DM population of glomeruli analyzed using OMP staining. However, we did find significantly reduced TAAR4+ OSN axon filling of the identified TAAR4 glomeruli 10 weeks after MMZ treatment (Mean TAAR4+ OSN fill, 63% for saline-treated vs. 18% for MMZ-treated; Fig. 5d), suggesting a deficit in reinnervation of TAAR4 glomeruli post-MMZ. Overall, these data suggest that TAAR4 glomeruli regenerate better than many other DM glomeruli, but still exhibit significant deficits in OSN axon reinnervation, in good agreement with our larger dataset for DM glomeruli (Fig. 4f).

**Figure 5.**
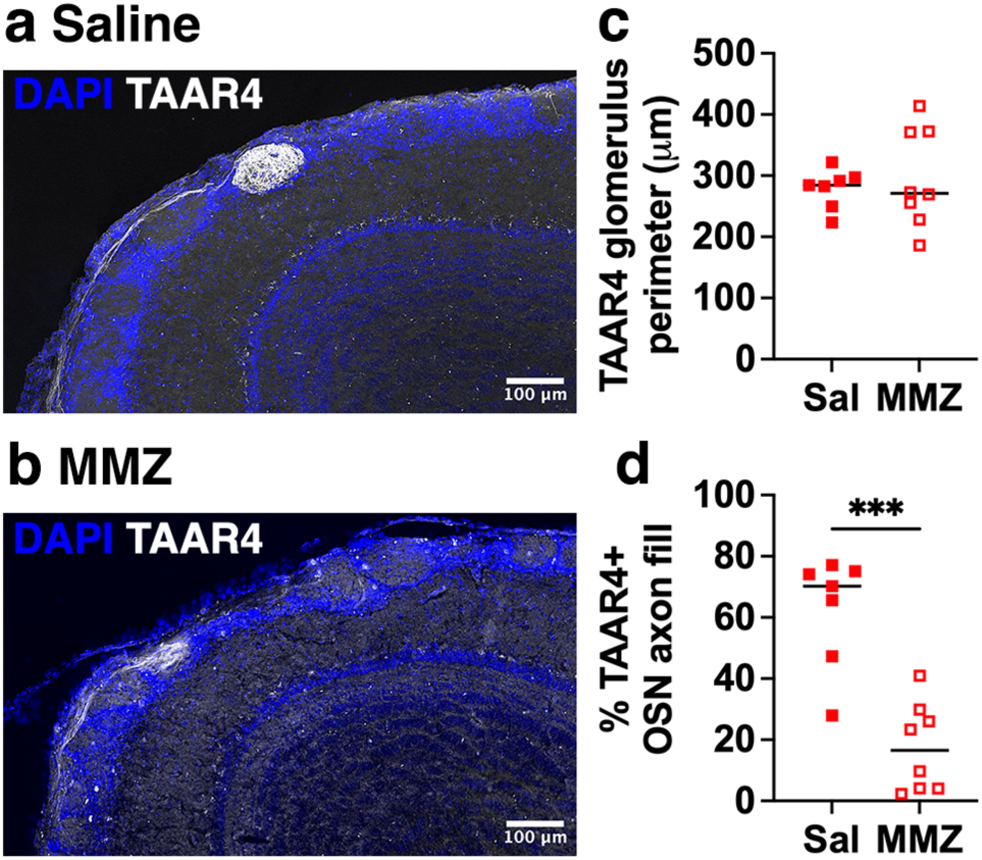
Reduced axonal filling of TAAR4 glomeruli 10 weeks after MMZ treatment. **a.** Example of a TAAR4 glomerulus in a saline-injected mouse. **b.** Example of a TAAR4 glomerulus 10 weeks after MMZ treatment. **c.** TAAR4 glomerulus perimeter is similar in saline- and MMZ-treated mice. P = 0.58, t = 0.57, n = 7 glomeruli from 3 saline-treated mice and 8 glomeruli from 3 MMZ-treated mice. **d.** Percentage TAAR4+ OSN axon fill of the TAAR4 glomerulus is significantly lower in MMZ-treated mice. P < 0.001, t = 5.23, n = 7 glomeruli from 3 saline-treated mice and 8 glomeruli from 3 MMZ-treated mice.

## Discussion

### Region-specific deficits in OB reinnervation after MMZ treatment

Regional differences in the extent to which newly generated OSN axons can reinnervate the OB have not previously been described. Using multiple different measures to assess the extent of recovery, we found marked deficits in reinnervation of dorsal OB glomeruli by new OSN axons that were particularly pronounced in the dorsomedial region. The most marked regional difference was in the percentage of glomeruli that was occupied by OMP+ axons, which was dramatically reduced only in the dorsomedial GL at 10 weeks post-MMZ. Our analysis of the TAAR4 glomerulus also found reduced reinnervation at 10 weeks post-MMZ. In contrast, two previous studies that used very similar measures of glomerular filling by mature OSN axons found recovery of this parameter within 28 days (after 75 mg/kg MMZ) [30] or 45 days (after 100 mg/kg MMZ) [29]. A key difference is that we analyzed all glomeruli in each coronal section and quantified the dorsomedial and dorsolateral regions separately, based on initial observations of differential recovery within the dorsal GL. In contrast, both previous studies [29,30] sampled a much smaller number of glomeruli distributed throughout the GL, so differences in dorsomedial glomeruli could have been missed or masked by averaging. In addition, differences in C57BL/6 mouse sub-strain or other experimental factors that alter the extent of OSN axon reinnervation could contribute to the discrepancy between our data and these previous studies [29,30].

Blanco-Hernández et al. used tagged OR lines to assess recovery of specific glomeruli after treatment with 75 mg/kg MMZ [37]. In line with our findings of extensive reinnervation of ventral and lateral glomeruli, Blanco-Hernández et al. found that projections of I7-expressing OSN axons to glomeruli in the ventrolateral OB recovered within 45 days of MMZ, although axons projected to two adjacent glomeruli, rather than the one glomerulus seen in untreated mice. They also found good recovery of M72-expressing OSN axons to the dorsolateral M72 glomerulus, albeit with many misrouted axons present even 90 days post-MMZ. This finding is not inconsistent with our data: while we found a reduced density of smaller glomeruli, there were many glomeruli present in the DL region at 10-20 weeks post-MMZ. Furthermore, the persistence of misrouted M72-expressing OSN axons supports our finding of deficits in reinnervation of dorsal surface glomeruli after MMZ treatment.

A previous study using MeBr treatment to ablate OSNs reported preferential reinnervation of the anterior OB compared to the posterior OB in rats [26]. In contrast, our data suggest no clear effect of position along the anterior-posterior axis or interaction between the effect of MMZ and position along the anterior-posterior axis on any of the parameters that were altered by MMZ treatment in at least one GL region. This discrepancy may be due to a combination of the method of OSN ablation and differences in reinnervation in rats vs. mice.

### Limited functional recovery of odor-evoked responses on the dorsal OB surface

Chronic imaging to track the extent to which neurogenesis restores odor-evoked responses following OSN cell death is currently only feasible for the dorsal surface of the OB. We found no recovery of odor-evoked responses in three of the six MMZ-treated mice that we imaged, and only very limited recovery in the other three, even up to 19 weeks after MMZ treatment. Collectively, less than 10% of the originally responding grid squares recovered a response to any odorant in the panel, and mean response amplitudes were an order of magnitude smaller than in the same fields of view imaged before MMZ treatment.

The only other study to use chronic imaging to assess functional recovery of odor-evoked responses in OSN axons used MeBr inhalation to ablate OSNs [38]. Cheung et al. found that synapto-pHluorin response magnitudes recovered to near control levels after 12 weeks, and that functional topography also recovered, with odorants evoking glomerular responses in similar regions to those before treatment. This contrasts with the poor functional recovery that we saw after MMZ treatment. However, Cheung et al. also found that glomeruli were smaller and had less well-defined borders, and that there were areas in which glomeruli were absent in some mice, in good agreement with our data. Why there was good recovery of odor-evoked synapto-pHluorin responses yet clear deficits in anatomical reinnervation in Cheung et al. is unclear. One factor that may be important is the variability in response to MeBr exposure across mice; indeed, only six out of ten MeBr-exposed mice were further analyzed in the Cheung et al. study, because the other four showed persistent odor-evoked responses four days after MeBr treatment, indicating a lack of OSN ablation. Another key factor in interpreting data from both MMZ and MeBr studies is recovery of the OE. MeBr can cause persistent loss of globose basal cells in the OE, resulting in areas that are not repopulated with OSNs, in a dose-dependent manner [26,38,52]. Not surprisingly, the extent of OB reinnervation by OSN axons correlates with OSN repopulation in the OE [26]. Hence, the incomplete OSN repopulation that was seen in Cheung et al. may account for deficits in glomerular reinnervation. In contrast, the OE can completely repopulate with OSNs after MMZ treatment, albeit taking longer (90d vs. 28d) when the dose is higher (100 mg/kg vs. 75 mg/kg) [29–31]. Hence, it is likely that the poor functional recovery of odor-evoked responses in the dorsal OB that we observed after MMZ treatment reflects a deficit in axonal reinnervation despite complete OSN repopulation. The mechanisms underlying this region-specific deficit in reinnervation will be important to elucidate in future studies.

Two other studies have provided alternative readouts of functional recovery after MMZ treatment. Browne et al. found rapid reconnection of OSNs with external tufted cells between 13 and 16 days post-MMZ [29]. They recorded from external tufted cells near medial and lateral glomeruli, which explains why they did not see the dorsal-specific deficits in OSN reinnervation that we observed. Blanco-Hernández et al. observed complete recovery of previously learnt odor discrimination tasks involving M72 and I7 receptor ligands, and partial recovery of innate aversion to TMT [37]. These mixed results likely reflect the anatomical location and extent of reinnervation of the glomeruli responsible for each behavior. Innate responses to TMT are mediated by both dorsal and ventral glomeruli [53,54], so the deficit in OSN reinnervation of the dorsal OB that we observed could underlie the partial recovery of TMT innate aversion seen by Blanco-Hernández. In contrast, Blanco-Hernández demonstrated significant reinnervation of the dorsolateral M72 glomerulus and the ventral I7 glomerulus, supporting the recovery of odor discrimination tasks employing their cognate ligands. It is also likely that behavioral recovery can occur even when glomeruli are only partially reinnervated, based on our previous studies at early time points post-MMZ in which mice could perform odor detection and discrimination tasks using only a small number of new, immature OSNs [48].

### Glomerular regeneration in the context of a developmental critical period

Regeneration of OB glomerular maps after OSN cell death requires accurate olfactory receptor-specific reinnervation of appropriate glomerular targets. During development, Glomerular maps become immutable after the closure of a perinatal critical period for OSN axon targeting [55,56]. Critical period closure correlates with cell death of a specialized population of navigator OSNs, which help ensure accurate targeting of OSN axons to appropriate glomeruli but are only present perinatally [57]. The absence of navigator OSNs in the adult OE raises the question of how accurate glomerular reinnervation might be achieved. Yet, multiple studies have shown substantial or complete recovery of glomerular targeting after OSN ablation [23,25,29,33,35,37,38,58]. One possible explanation is that degenerating OSN axons provide a pathway for glomerular targeting via homotypic interactions between odorant receptors [26,56,57]. This seems plausible given that degenerating axons are present for at least two weeks after MMZ treatment [29,30], and the first newborn OSNs begin to reinnervate the OB as early as five days post-MMZ [48]. However, it leaves open the question of why there is a selective deficit in reinnervation of dorsal, and particularly dorsomedial, glomeruli after MMZ treatment. Region specific differences in the number of projecting OSN axons, or the rates of OSN axon degeneration vs. arrival of new OSN axons could play a role. In addition, axon guidance mechanisms that specify projections along the dorsal-ventral axis of the OB could be responsible. Both Slit-Robo and semaphorin-neuropilin expression gradients play important roles in dorsoventral patterning of OSN axonal projections [59–62]. However, the role of these axon guidance molecules during large-scale regeneration is not clear. For example, during development, OSNs projecting to the dorsal OB are generated early, and Sema3F secretion by these axons is repulsive to axons of later-arriving Nrp2-expressing OSN axons that therefore project to the ventral OB [62,63]. However, in this study, we saw good reinnervation of the ventral but not the dorsal OB, suggesting that different axon guidance mechanisms may be invoked following large-scale OSN ablation in adults.

Another potential explanation might involve differences in OSNs themselves. Two different classes of OSNs that project to the dorsal OB, which are strongly biased to express class I vs. class II ORs, have previously been described, with class I OR-expressing OSNs projecting predominantly to the anterodorsal and dorsomedial OB [53,64,65]. Interestingly, class I OSNs may either be injured preferentially or regenerate less well in post-viral olfactory dysfunction patients [66], although the mechanism underlying this is unclear. A specific subset of dorsomedial projecting OSNs is also evident in P-LacZ-Tg mice, in which β-gal expressing OSN axons project to many glomeruli throughout the OB, but avoid the dorsomedial region [64]. Furthermore, OSNs in zone 1 of the OE, which corresponds to the dorsomedial region of the OE, selectively express the enzyme NADPH:quinone oxidoreductase (NQO1), which catalyzes the reduction of quinones [67]. Interestingly, OMP staining is selectively reduced in dorsomedial NQO1+ glomeruli after 10 months (but not after 4 months) on a protein-restricted diet [68]. Long-term protein-restricted mice had fewer mature OSNs in zone 1 of the OE, as marked by NQO1 expression in OSNs. There was also evidence of NQO1-mediated oxidative stress in zone 1 OE cells, and expression of apoptotic markers in those same cells, which may underlie the reduced density of mature OSNs. Another study found that the density of both proliferating cells and mature OMP+ OSNs was lower in the NQO1+ OE vs. the NQO1-OE two weeks post-MMZ, an effect that was further exacerbated by sleep deprivation [69]. The authors suggested this may be due to the previously reported reduced bioavailability of retinoic acid (RA) in HBCs within the NQO1+ OE due to cytochrome P450 family 26-mediated RA degradation [70], which may result in reduced injury-induced HBC proliferation. It is therefore possible that a similar mechanism could underlie the strong reduction in glomerular re-innervation by OMP+ OSNs 10 weeks post-MMZ in our study. Elucidating the mechanisms that mediate successful vs. unsuccessful reinnervation of OSN axons may have clinical ramifications for the treatment of anosmia, as well as broader implications for therapeutic strategies to promote axon regeneration elsewhere in the nervous system.

## Acknowledgments

Research reported in this manuscript was supported by the National Institutes on Communication Disorders and Stroke (National Institutes of Health) R01DC018516 to C.E.J.C. and F32DC018431 to T.K. The content is solely the responsibility of the authors and does not necessarily represent the official views of the National Institutes of Health. We thank Dr. Thomas Bozza for providing the TAAR4-RFP and TAAR4-ChR2-YFP mice, and members of the Cheetham lab for helpful discussions.

## Supplementary Material

**Table S1.**
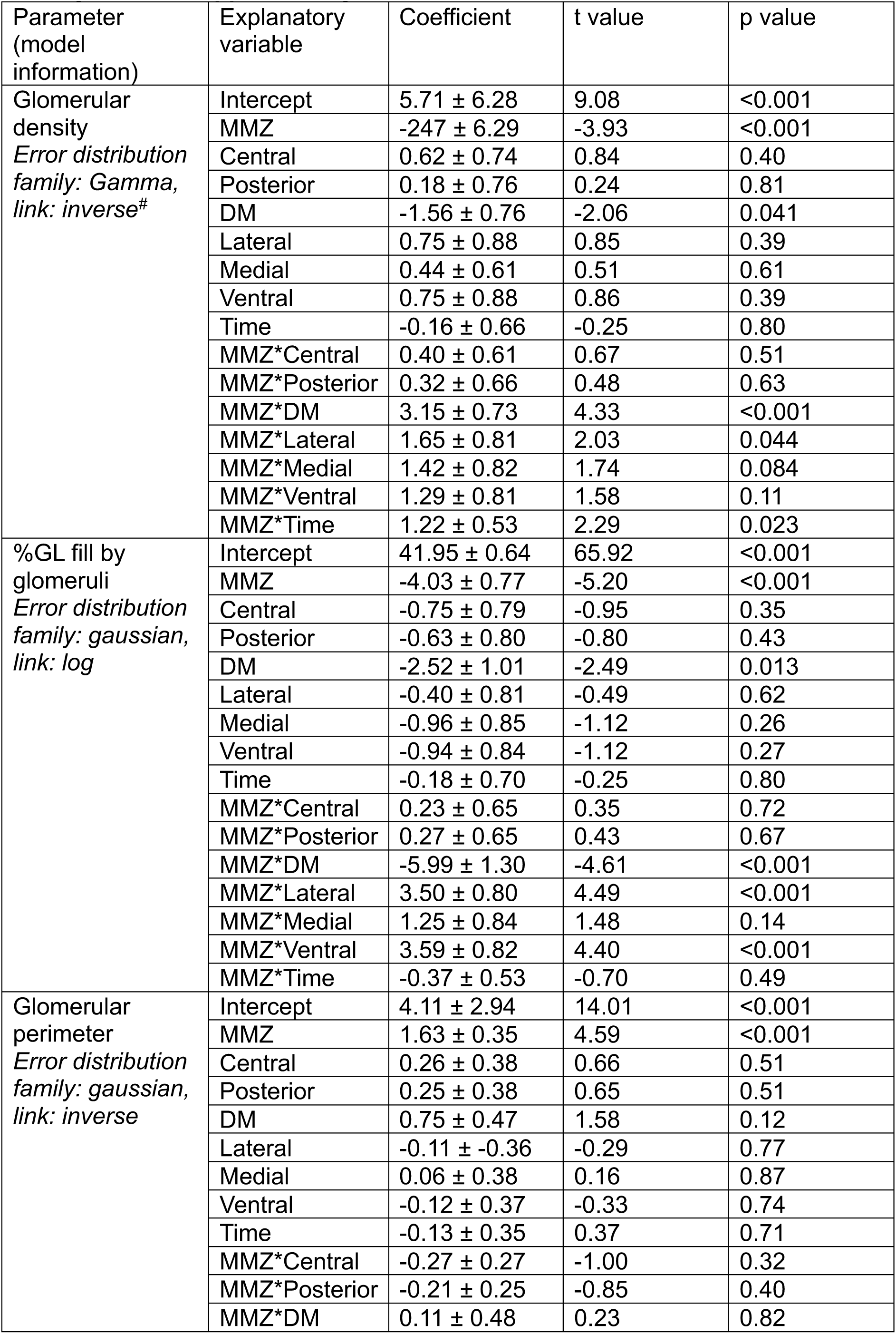
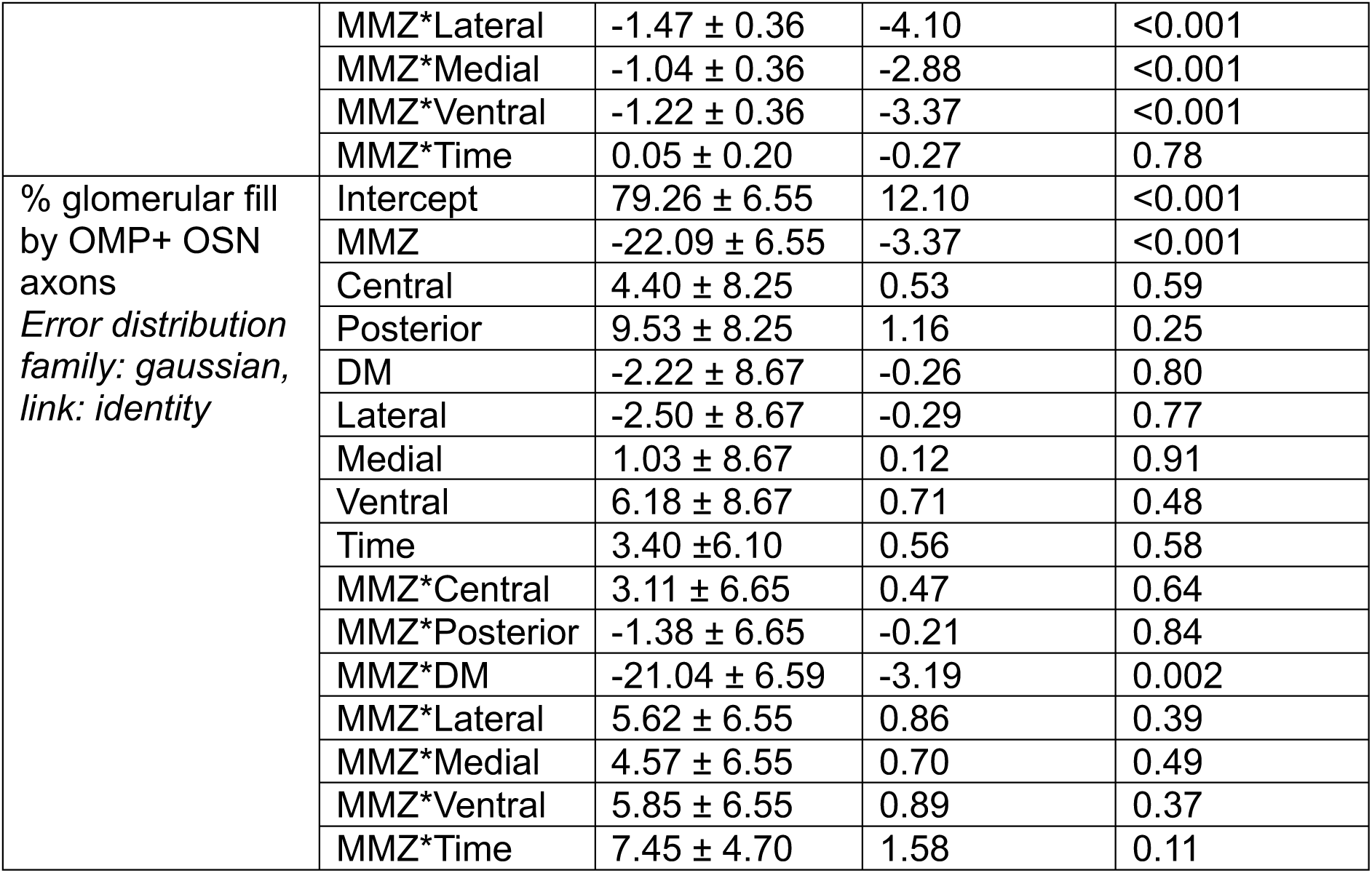
No effect of position along the anterior-posterior axis on parameters that showed deficits in regeneration post-MMZ. GLMs were used to assess main effects and interactions with MMZ treatment. Model variables were as follows: MMZ, mice treated with MMZ; Central, OB section ∼50% along the anterior-posterior axis; Posterior, OB section ∼75% along the anterior-posterior axis; DM, dorsomedial GL; Lateral, lateral GL; medial, medial GL; ventral, ventral GL, Time, mice at 20 weeks post-MMZ or age-matched controls. For analysis of glomerular density, coefficients are values ×10^−3^. For analysis of % GL fill by glomeruli, coefficient values are ×10^1^. For analysis of glomerular perimeter, coefficients are values ×10^−3^. For analysis of % glomerular fill by OMP+ OSN axons, coefficient values are as stated.

